# Sequence- and structure-specific cytosine-5 mRNA methylation by NSUN6

**DOI:** 10.1101/2020.10.01.320036

**Authors:** Tommaso Selmi, Shobbir Hussain, Sabine Dietmann, Matthias Heiss, Jean-Michel Carter, Rebecca Dennison, Sophia Flad, Ya-Lin Huang, Stefanie Kellner, Susanne Bornelöv, Michaela Frye

**Author notes:** Corresponding authors: SB MF. Equal contribution (Tommaso Selmi, Shobbir Hussain).

## Abstract

Methylation is the most common internal modification in mRNA. While the highly abundant N6-methyladonsine (m^6^A) modification affects most aspects of mRNA function, the precise functions of the rarer 5-methylcytosine (m^5^C) remains largely unknown. Here, we map m^5^C in the human transcriptome using methylation-dependent individual-nucleotide resolution cross-linking and immunoprecipitation (miCLIP) combined with RNA bisulfite sequencing. We identify NSUN6 as a methyltransferase with strong substrate specificity towards mRNA. NSUN6 primarily targeted three prime untranslated regions (3’UTR) at the consensus sequence motif CTCCA, located in loops of hairpin structures. Knockout and rescue experiments revealed that only mRNA methylation sites containing the consensus motif depended on the presence of NSUN6. Furthermore, ribosome profiling demonstrated that NSUN6-specific consensus motifs marked translation termination. However, even though NSUN6-methylated mRNAs were reduced in NSUN6 knockout cells, NSUN6 was dispensable for mouse embryonic development. Thus, our study identifies NSUN6 as methyltransferase targeting mRNA in a sequence- and structure-specific manner.

## INTRODUCTION

The over 170 known RNA modifications extensively increase the functional diversity of RNA molecules (1). Accordingly, RNA modifications emerged as an important additional regulatory layer of gene expression programs and are often required for normal development (2). Recent advances in detection technology, mostly associated with high-throughput sequencing, revealed how RNA modifications influence many stages of RNA metabolism, and thereby effect diverse biological processes including cell fate decisions, immune responses, and tumorigenesis (3-5).

One of the best-studied modifications is N6-methyladenosine (m^6^A), the most abundant mRNA modification that controls gene expression (6). The prevalence and function of other rarer mRNA modifications such as N1-methyladenosine (m^1^A), N6,2’-O-dimethyladenosine (m^6^Am), m^5^C, 5-hydroxymethylcytosine (hm^5^C), and Pseudouridine (Ψ) are less well-characterised and often somewhat controversial (7). One of the most disputed modifications in mRNA is m^5^C as current conventional detection methods are associated with high background levels, making this low-abundance modification particularly challenging to define (8,9). Furthermore, the total levels of m^5^C in mRNA varies between tissues (10). Nevertheless, NSUN2, a m^5^C RNA methyltransferase mainly targeting tRNAs, has consistently been linked to mRNA methylation (11-15).

NSUN2 is one of eight evolutionary conserved m^5^C RNA methylases (NSUN1-7 and DNMT2) (2). Of these, NSUN2, 3, 6 and DNMT2 have all been shown to methylate tRNAs, yet in a non-overlapping and site-specific manner (15-20). NSUN2 is however the only enzyme with a broader substrate-specificity, methylating the majority of expressed tRNA, other abundant non-coding RNAs and a small number of mRNAs (11-15,21).

Here, we map NSUN6-dependent m^5^C sites in RNAs in the human transcriptome using our recently developed miCLIP method (12,22). Unexpectedly, we find that most sites located to coding RNAs within the consensus sequence motif CTCCA. NSUN6-specific sites were enriched in the 3’UTR and marked translation termination. RNA bisulfite (BS) sequencing confirmed m^5^C sites in mRNAs that were lost in knockout cells and rescued by over-expressing the NSUN6 protein. Although mRNAs stability was enhanced in the presence of NSUN6-mediated m^5^C, NSUN6 was dispensable for mouse embryonic development. Our study shows for the first time that NSUN6 mediates site-specific deposition of m^5^C in mRNA.

## MATERIAL AND METHODS

### Cell culture

The human embryonic stem cell (hESC) line Hues9 (H9) was obtained from the Wicell Research Institute (Madison, WI) and maintained in Essential 8 media (Thermo Fisher Scientific) on hESC-Qualified Matrigel (Corning) coated plates. Media was refreshed daily and the cultures were dissociated in clumps every 4 days using 0.5mM EDTA in PBS. In the embryoid bodies experiments, 70% confluent hESC were dissociated in clumps and seeded on ultra-low attachment well plates (Corning) maintaining a 1:1 dilution factor. After plating, hESC were cultured in Essential 6 media (Thermo Fisher Scientific) plus 10 μM rock-inhibitor (Y-27632) (Stem Cell Technologies Canada) for the first 24 hours and for further 4 to 6 days in Essential 6 media (Thermo Fisher Scientific).

HEK293 (HEK) cells were grown in DMEM Media (Thermo Fisher Scientific) supplemented with 1mM Glutamax (Thermo Fisher Scientific), 10% heat inactivated FBS (Thermo Fisher Scientific). All cells were grown at 37°C, 5 % CO2.

### Generation of NSUN6 knockout, rescue and overexpressing lines

NSUN6 knockout H9 cells were generated either by using homology directed recombination (HDR) or by inserting random Indels in response of double strand breaks (NHEJ). For HDR, we targeted exon 2 of NSUN6 with wild type Cas9 (pSpCAs9(BB)2A-GFP) plus the recombination vector pD07 (Genecopoeia) carrying the selection genes puromycin and eGFP under control of EIF1a promoter and with homology arms on Introns 1 and 2. gRNAs were designed using the gRNA design software from the Feng Zhang lab at MIT (Exon2 gRNA1: ATT TTT CAC ATG TTG TAC TG **AGG**, Exon2 gRNA2: GAT GAA CTT CAG AAG GTT TG **TGG)**. 4 days after nucleofection with the AMAXA Nucleofector Kit (Lonza), we applied puromycin selection until we observed the appearance of green colonies, which were screened for integration of the recombination cassette and for the presence of NSUN6 exon 2. For NHEJ, we targeted the exon 9 of NSUN6 with wild type Cas9 (pSpCAs9(BB)2A-GFP) (Exon 9 gRNA: ATC CAG AAG AAT TCG GTC AA **AGG**), 3 days after nucleofection, the targeted cells were sorted and re-plated at low density in E8 conditioned media. We then screened by Sanger Sequencing the grown colonies for InDels in NSUN6 exon 9. For generating NSUN6 overexpressing H9, NSUN6 CDS was PCR amplified from the TrueORF pCMV-Entry vector (Origene) and cloned via Gibson assembly (NEB) into the destination piggybac vector pBPCAG-cHA-IN (kindly provided by Austin Smith). This vector and the piggy bac transposase were then nucleofected into H9 using the AMAXA nucleofector kit (Lonza). Puromycin selection started 4 days following nucleofection and the surviving clones were screened by qPCR and Western Blot for NSUN6 expression. The same overexpression strategy used in H9 was also adopted to rescue NSUN6 expression in knockout HEK293 cells.

### Mass spectrometry

NSUN2, NSUN6 and DNMT2 were depleted by transfecting the corresponding siRNA Pools or a Negative Control siRNA pool (siTOOLsBIOTECH). The siRNAs were designed to target the transcripts coding for NSUN2 (NCBI Gene ID: 54888), NSUN6 (NCBI Gene ID: 221078) and DNMT2 (NCBI Gene ID: 1787). For sample collection, cells were directly lysed in Trizol (ThermoFisher Scientific) and total RNA was extracted according to the manufacturer’s instruction. Mass spectrometry to quantify m^5^C was performed as described previously (10,23).

### Generation of the Nsun6 knockout mice

All mice were housed in the Wellcome Trust-Medical Research Council Cambridge Stem Cell Institute Animal Unit. All mouse husbandry and experiments were carried out according to the local ethics committee under the terms of a UK Home Office license P36B3A804 and PPL70/7822.

Two embryonic stem cell lines containing a knockout first allele (with conditional potential) were obtained from EuMMCR (*Nsun6* ^tm1a(EUCOMM)Hmgu^). Mice homozygote for the targeted trap allele was used to analyse *Nsun6* total knockout. Genotyping primers were: *Nsun6*-forward (AAT CCA GCA TTC CTG TTG TTC AGC), LoxR (TGA ACT GAT GGC GAG CTC AGA CC), *Nsun6*-5’arm-2 (ACA GTG AGT CAG GTG AGG TGT GCC), and *Nsun6*-rev (CAC AAT GAG ACA GCA CCC AG). The LacZ-neo cassette was used as reporter for *Nsun6* RNA expression in wild-type and *Nsun6* total knockout mice. LacZ staining on whole mounts and sections of embryos was performed as described previously (24).

### RT-qPCR and Western blotting

Total RNA was extracted using TRIZOL (Thermo Fisher Scientific) according to manufacturer instructions. Reverse transcription was performed using SuperScript III Reverse Transcriptase (Thermo Fisher Scientific) and random primers (Promega). Quantitative PCR were run using TaqMan probes (Thermo Fisher Scientific) for eukaryotic 18S rRNA (X03205.1), MARCKSL1 (Hs00702769_s1), TRAF7 (Hs00260228_m1), ANGEL1 (Hs00380490_m1), CALM3 (Hs00968732_g1), BAG6 (Hs00190383_m1), CUX1 (Hs00738851_m1), TRIMM50 (Hs01390531_m1), BUB3 (Hs00945687_m1), EEF (Hs00265885_g1), MACF1 (Hs00201468_m1), DLX5 (Hs01573641_mH), DNMT3B (Hs00171876_m1), FOXD3 (Hs00255287_s1), GATA6 (Hs00232018_m1), HOXA1 (Hs00939046_m1), NANOG (Hs02387400_g1), POU5F1 (Hs03005111_g1), TDGF1 (Hs02339497_g1).

For protein isolation, cells were first rinsed with PBS and lysed in ice-cold RIPA buffer (50mM Tris-HCL pH 7.4, 1% NP-40, 150mM NaCl, 0.1% SDS, 0.5% Sodium deoxycholate). RIPA was supplemented with cOmplete Mini EDTA-free Protease Inhibitor Cocktail tablets (11836170001, Roche). Cells were collected using a cell scraper and the lysates were centrifuged for 15 minutes at maximum speed in a pre-cooled centrifuge at 4°C, and their supernatant collected and kept on ice. Cell protein lysates were mixed with NuPAGE LDS Sample Buffer (4X) (NP0007; Invitrogen) and run on polyacrylamide gels. Proteins were transferred to a nitrocellulose or PVDF membrane (GE Healthcare). Membranes were blocked for a minimum of 1 hour at room temperature in 5% (w/v) non-fat milk or 5% (w/v) BSA (A4503-50G; Sigma Aldrich) in TBS-T (1x TBS and 0.1% Tween-20) and then incubated with primary antibody in blocking solution overnight at 4°C. Each membrane was washed three times for 10 minutes in TBS-T prior to incubation with the appropriate Horseradish peroxidase (HRP)-labeled secondary antibody (1:10 000) in TBS-T at room temperature for 1 hour. After washing, the antibodies were detected by using the Amersham ECL Prime Western Blotting Detection Reagent (RPN2232; GE Healthcare). The primary antibody was NSUN6 (1:500, 17240-1-AP, Proteintech). Anti-γ-Tubulin (T6557; Sigma Aldrich) or Ponceau staining served as a loading control.

### miCLIP

Full-length cDNA constructs for NSUN6 in the pCMV6-Entry-Myc vector were obtained from OriGene. Site-directed mutagenesis to generate the miCLIP-mutants was performed using the QuikChange II Site-Directed Mutagenesis Kit from Agilent as per the manufacturer’s instructions. To generate the NSUN6 miCLIP mutant, cysteine 326 was mutated to alanine using the following primers: (forward) GAA TTC TTC TGG ATG CAC CCG CTA GTG GAA TGG GAC AGA GAC; (reverse) GTC TCT GTC CCA TTC CAC TAG CGG GTG CAT CCA GAA GAA TTC. HEK293 cells were transfected with either NSUN6 wild-type and the miCLIP-mutant construct using Lipofectamine 2000 (Life Technologies) and harvested 24 hours later.

The harvested cells were lysed in lysis buffer consisting of 50 mM Tris-HCL pH 7.4, 100 mM NaCl, 1% NP-40, 0.1% SDS, 0.5 % sodium deoxycholate. Lysates were then treated with high concentration of DNase and low concentration of RNaseI to partially fragment RNAs. Lysates were cleared by centrifugation at 13,000 rpm for 15 minutes at 40°C and then incubated with Protein G Dynabeads (Life Technologies) in the presence of an anti-Myc antibody (9E10, Sigma). Following stringent washing, 3’ end dephosphorylation was performed with T4 polynucleotide kinase (New England Biolabs) before addition of a pre-adenylated linker using RNA ligase (New England Biolabs). 5’ end labelling was then performed using T4 PNK and ^32^P-ATP before protein-RNA complexes were eluted and run on denaturing gels. Nitrocellulose transfer was performed, and the radioactive signal was used to dissect nitrocellulose pieces that contained NSUN6-partially digested RNA complexes. RNA was recovered by incubating the nitrocellulose pieces in a buffer containing Proteinase K and 3.5 M urea. Next, reverse transcription was performed using oligonucleotides containing two inversely oriented adaptor regions separated by a BamHI restriction site. cDNAs were size-purified on TBE-Urea gels before being circularized by CircLigase II (Epicentre). Circularised cDNAs were then annealed to an oligonucleotide complementary to the BamHI site and then BamHI digested. Linearized cDNAs were then PCR-amplified using primers complementary to the adaptor regions using 25 cycles of PCR. Libraries were then subjected to high-throughput sequencing using the Illumina HiSeq 2000 platform.

### RNA bisulfite conversion and sequencing

Total RNA from H9 wild-type (WT), knockout (KO), overexpression (OEX) and HEK293 WT, KO and NSUN6 rescue (RES) cells was first extracted using Trizol (Thermo Fisher Scientific) and subsequently treated withDNaseI (Ambion) and RiboZero (Illumina) to remove contaminating DNA and ribosomal RNAs. The remaining RNA was then converted as previously described (15,25). Briefly, 10 μg of RNA was resuspended in 10 μl of RNAse free water and mixed with sodium bisulfite pH 5.0 (42.5 μl) and DNA protection buffer (17.5 μL) (EpiTect Bisulfite Kit, Qiagen). The deamination reaction was then carried out by incubating in a thermal cycler for four cycles of 5 minutes at 70°C followed by 1 hour at 60°C and then desalted with Micro Bio-spin 6 chromatography columns (Bio-Rad). RNA was desulphonated by adding an equal volume of 1 M Tris (pH 9.0) to the reaction mixture for 1 hour at 37°C, followed by ethanol precipitation. The bisulfite-converted RNA quality and concentration were assessed on a Bioanalyzer 2100 RNA nano-chip (Agilent). About 120 ng of bisulfite-converted RNA were used to generate Bisulfite-seq libraries using the TruSeq Small RNA preparation kit (Illumina). Before library preparation, the fragmented RNA wasend-repaired with T4 PNK and Spermidine (New England Biolabs). The size selection step was omitted, as the bisulfite-converted RNA was sufficiently fragmented by the previous conversion reaction. First the Illumina RNA adapters were then ligated, reverse-transcribed at 50°C for 1 h ourwith SuperScript III and 1 mM of each dNTP (SuperScript III cDNA synthesis kit, Invitrogen) followed by 18-cycle PCR amplification.

### RNA sequencing and Ribosomal Profiling

Total RNA extraction, Ribosmal profiling and libraries for H9 NSUN6 WT and KO cells (at least 4 replicates each) were performed as described before (26-28). The samples were multiplexed and sequenced on the HiSeq 4000 platform (Illumina).

### Processing, mapping and quantification of RNA-seq and Ribo-seq reads

The H9 RNA-seq data was sequenced as paired-end 2×150 nt and the HEK RNA-seq data was sequenced as single-end 50 nt. All Ribo-seq data was single-end 50 nt. To process the data, Trim galore! (https://github.com/FelixKrueger/TrimGalore) with parameters “--stringency 6 -e 0.1” and “--paired” for the paired-end data was first used to remove Illumina adapters and to exclude trimmed reads shorter than 20 nt. Alignment was done using Tophat2 (v2.1.0) using an index with known transcripts (Gencode v23) as guidance and with novel splice junctions were permitted. The RNA-seq reads were aligned directly to the reference genome (hg38). Ribo-seq reads were firstly aligned to a set of known rRNA and tRNAs (selected from the UCSC RepeatMasker tracks), followed by alignment of all unmapped reads to the reference genome. Multi-mapping read were excluded. FeatureCounts was used to quantify the number of reads per gene using the Gencode v23 gene models. Only reads aligning to the sense strand of the gene, represented either by its exons (RNA-seq) or its coding sequence (Ribo-seq), and with mapping quality at least 20 were counted. For the paired-end RNA-seq the additional flags “-p -B -C” was specified to exclude chimeric reads and/or reads mapping with only one end.

### RNA-seq differential expression analyses

Differential expression analysis was done using the R Bioconductor edgeR package. Genes with mean expression below 1 counts per million were considered non-expressed and excluded from the analysis. The R Bioconductor cqn packages (29) was used for conditional quantile normalization to calculate offsets correcting for gene lengths (from featureCounts) and GC content (from biomaRt). The offsets were passed to the edgeR DGEList object, followed by a likelihood ratio test using glmFit and glmLRT. Five knockout cell lines (four H9 and two HEK) were compared to their respective control cell line (two H9 and one HEK). Each cell line was sequenced in four replicates.

### Determining read periodicity and codon enrichments using Ribo-seq

Codon usage in the Ribo-seq data was calculated following (28). The analysis was focused on reads of length 27-29, which were showing the strongest periodicity. Position 12-14 were determined to correspond to the codon at the P-site.

The bam file for each sample with uniquely aligned reads was converted to bed format. Bedtools intersect was used to select reads with at least 50% overlap to Gencode-annotated coding sequences. Next, the reading frame of the 5’ end of each read was determined using the frame information in the Gencode annotation. If the frame did not agree with the expected reading frame for that read length, the read was discarded. Then, nucleotide positions 1-27 were extracted from each read as nine codons, numbered as codon position -5 to +3, where 0 corresponded to the A-site. As expected, position -4 to -2 and +1 to +3 correlated well to the genome-wide distribution of codons in the human translatome, whereas counts from the predicted P-site and A-site did not.

The number of codon occurrences were counted separately for each ribosome-protected codon position and converted into a fraction of the total number of codons. Normalized codon counts were obtained by dividing the codon fraction at a specific position by the mean fraction across all nine positions.

### Processing and mapping of miCLIP reads

In order to reduce amplification bias, the primers used for reverse transcription during miCLIP experiments were designed to include a 6-nucleotide random barcode at positions 1-3 and 8-10 to enable tracing of individual cDNAs. Reads were de-multiplexed using the experimental barcode at positions 4-7, and reads with identical random barcodes, representing PCR products, were filtered. The number of different random barcodes for each unique read, which represented cDNA counts, was stored for further analysis. Barcodes were trimmed from the 5’end, and the adapter sequence ‘AGATCGGAAGAGCGGTTCAG’ from the 3’end of the reads with *cutadapt* (https://code.google.com/p/cutadapt; options: ‘-O 4 –e 0.06’), and only reads with a minimal length of 18 nt were retained.

Trimmed miCLIP reads were mapped to the human reference genome (UCSC GRCh37/hg19) by using *bowtie* (http://bowtie-bio.sourceforge.net/index.shtml) with parameters *‘m 1 v 1 best strata*’ to select uniquely mapping reads allowing one mismatch. Methylation sites were thus inferred from miCLIP read truncation positions by assigning the read counts to the closest cytosine within +/-2nt of the truncation site. Pooled read counts per cytosine were normalized per million uniquely mapping reads (RPM). If not stated otherwise, only high-confidence methylation sites with normalized read counts > 50 RPM in at least two out of three replicates were selected for downstream analyses.

### Quantifying coding sequence annotation and Ribo-seq/RNA-seq reads at miCLIP sites

To visualize coding sequence across miCLIP sites, annotated coding sequences (Gencode V28) were merged using bedtools merge and was then converted to bigWig format using the UCSC bedGraphToBigWig tool. This was done separately for both strands. Furthermore, to allow visualization of RNA-seq and Ribo-seq read coverage at miCLIP sites, deepTools bamCoverage was used to convert the aligned reads to bigWig files. Each strand was quantified separately, and a blacklist file containing all rRNA, tRNA, snoRNA, snRNA, and miRNA regions was provided. The bin size was set to 1 and an offset of 12 was used to only consider a single nucleotide corresponding to the “P” site predicted from each read.

Next, deepTools computeMatrix in “reference-point” mode with parameters -b 1500 -a 1500 --missingDataAsZero” was used to extract annotation or reads coverage across miCLIP sites. To classify the miCLIP sites as start, middle and end of translation the difference between the number of covered bases upstream and downstream of the miCLIP site was used. The standard error (SE) of the differences was calculated and a threshold of +/-1.96SE was used to defined ‘start’ and ‘end’ of translation. The remaining sites were classified as ‘middle’. Custom R scripts were used to combine sites on both strands and to visualize it as a heatmap or a profile plot. The scripts related to this analysis are available at https://github.com/susbo/Selmi-et-al-scripts.

### Calculating overlap between miCLIP sites and differentially expressed genes

Out of the 15,885 genes expressed in HEK, 1,853 genes were defined as down-regulated and 2,008 as up-regulated at padj < 0.05 and abs(log2FC) > 0.5 in both HEK knockout clones. Next annovar (30) was used to identify the closest genes for all 252,135 putative miCLIP sites. The set of genes with exonic miCLIP sites (at 0, 0.5, 1, 3, 5, 10, or 50 cpm) was compared with genes without exonic miCLIP sites. The relative enrichment or depletion of gene down-or up-regulation was calculated as an odds ratio with a 95% confidence interval using Fisher’s exact test.

### Processing and mapping of BS-seq reads

The BS-seq was processed as described previously (21). First, Trim Galore! (v0.4.0) with parameters “--stringency 3 -e 0.2 -a TGGAATTCTCGGGTGCCAAGGA” was used to remove sequencing adapters and exclude reads shorter than 20 nucleotides. Next, alignment to the hg38 reference genome was done using Bismark (v0.14.4) with parameters “-n 2 -l 50 --un --ambiguous --bowtie1 --chunkmbs 2048”, to allow for up to two mismatches and to save unaligned and ambiguously mapping reads separately. Seqtk with parameters “-e 3” was used to remove the last three bases (a potential tRNA ‘CCA’ tail) from the unaligned and ambiguous reads followed by a second alignment attempt using Bismark. Finally, ngsutils (v0.5.9) in the “junction” mode was used to extract splice junctions from known genes (Gencode v28) and unaligned reads from the second alignment were aligned to the junctions using Bismark. Reads aligned to the junctions were converted back to genomic coordinates using bamutil (https://github.com/statgen/bamUtil) in “convertregion” mode. ‘N’ in the cigar string was replaced with ‘D’ for compatibility with bismark_methylation_extractor.

Samtools merge was used to combine aligned reads from all three alignment attempts. Reads with >1/3 methylated cytosines were discarded as they are likely artifacts from conversion-resistant regions. The bismark_methylation_extractor with the “—bedGraph --counts --CX_context” options was used to extract methylated cytosines.

The BS-seq data are available on GEO with the accession number GSE125046. miCLIP data are available under the accession number GSE66011.

For comparisons with methylation sites in Huang et al. (11) we used the published methylation sites from their Supplementary Table 4. The liftover tools was used to convert hg19 coordinates to hg38 coordinates. However, to visualize the methylation levels in those samples, we re-aligned those datasets to the hg38 genome using our pipeline.

### Quantification of methylation at miCLIP sites and known methylation sites

Methylation signal was quantified across protein-coding miCLIP sites and across the 13 published lists of methylation sites from Huang et al. (11) representing HEK293, HeLa (control and NSUN2 knockout) and seven tissues. Genomic coordinates were converted to hg38 using the liftover tool, and the genomic context (+/-10 nucleotides) of the site was extracted. For each set of sites, we analyzed sites with the CTC[CT]A motif and sites without the motif separately. To reduce the noise from the miCLIP experiment, miCLIP sites located next to a motif were shifted to the motif and duplicate sites were removed. For each cytosine within 10 nucleotides from the methylation site, the total number of reads that either supported or did not support methylation were calculated. To avoid having very high-covered sites dominating the analysis, sites with more than 5 reads were normalized to 5 reads before the reads were summarized. Methylation level was calculated as the number of normalized reads supporting methylation, divided by the total number of normalized reads.

## RESULTS

### miCLIP reveals substrate specificity for NSUN6

RNA:m^5^C methyltransferases contain a catalytic domain with a common structural core and the S-Adenosyl methionine-binding site (Figure S1A). Two conserved cysteines, both located within the methyltransferases active site, are required for completion of the catalytic process (31). Methylation is initiated when cysteine (C1) forms a covalent bond with the cytosine pyrimidine ring (32). The second conserved cysteine (C2) is required to then break the covalent adduct and to release the methylated RNA (Figure S1B). Mutating C2 to alanine results in the irreversible formation of an enzyme-RNA crosslinked complex precisely at the already methylated cytosine (Figure S1C) (31,33,34). We previously utilized this irreversible formation of catalytically-crosslinked complexes to identify NSUN2 and NSUN3 methylated nucleosides genome-wide via miCLIP (Figure 1A; Figure S1A-C) (12,17). Here, we analysed a third related m^5^C RNA methylase, NSUN6. We sequenced all RNAs crosslinked to the mutated NSUN6 protein (Figure 1B) and found that the majority of miCLIP sites located to mRNAs (Figure 1C; Table S1). In contrast to NSUN6, NSUN2-specific miCLIP sites mainly occur in tRNAs (Figure 1D; Table S2) (12). Thus, NSUN6 has a distinct substrate specificity, different from NSUN2.

**Figure 1.**
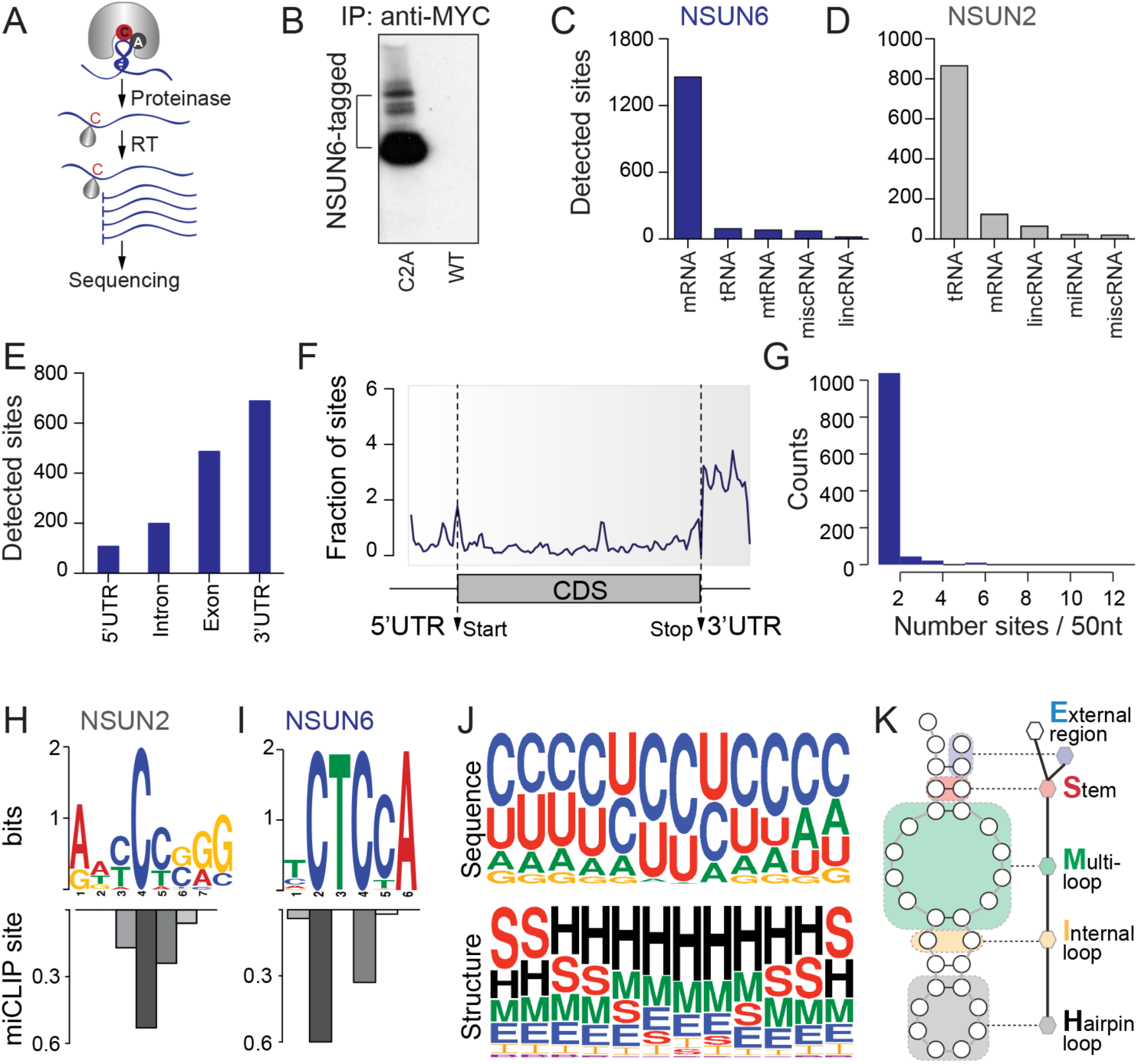
NSUN6 miCLIP reveals consensus motif in targeted RNA. (A) Schematic overview of the miCLIP method. (B) Polyacrylamide gel showing NSUN6-tagged proteins and released RNA after RNase treatment, which was isolated and sequenced. C2A: construct carrying the point mutations C->A; WT: wild-type construct. (C, D) Number of NSUN6 (C) and NSUN2 (D) miCLIP sites in the indicated RNAs. (E) Number of NSUN6 miCLIP sites in protein coding RNAs. (F) Distribution of NSUN6 miCLIP sites along mRNA. (G) Number of detected NSUN6 miCLIP sites within 50 nucleotides windows. (H, I) Binding motifs (upper panels) of NSUN2 (H) and NSUN6 (I) and frequency of miCLIP sites at the respective position (lower panels). (J, K) GraphProt identified sequence (J; upper panel) and structural motif (J; lower panel) of NSUN6-targeted sites and predicting of the methylated sites in hairpin (H) loops of stem loop structures (K).

### NSUN6 targets 3’ UTRs in a sequence- and structure-specific manner

We previously demonstrated that the small number of NSUN2-specific miCLIP sites in mRNAs mostly occurred in exons of nuclear-encoded genes (12). In contrast, NSUN6-mediated methylation occurred mainly along the 3’UTR (Figure 1E, F). The advantage of miCLIP over other RNA immunoprecipitation (RIP) methods is the covalent cross-link of the enzyme to the cytosine undergoing methylation, allowing the detection of the methylation site at nucleotide-resolution (12,22). Accordingly, we found that NSUN6-specific target sites predominantly occurred as single sites (Figure 1G). In addition, we found that the methylated sites occurred in a sequence-specific manner. NSUN6 targeted a clear consensus sequence, and this motif was present in the vast majority (80%) of all targeted sites (Figure 1H, I; Figure S1D). Since RNA secondary structures can modulate protein binding (35), we next asked whether local RNA structure might influence NSUN6 targeting. We determined the sequence- and structure-specific preferences of NSUN6 miCLIP sites using GraphProt (36), and found that the methylated sites occurred preferentially within hairpin loops (H) of stem loop structures (Figure 1J, K). In conclusion, methylation by NSUN6 is based on a defined sequence-structure element in mRNAs.

### RNA bisulfite sequencing confirms NSUN6-specific methylation sites

The miCLIP method relies on over-expressing the mutated protein (37). To confirm methylation sites in mRNA deposited by the endogenous NSUN6 protein, we performed RNA bisulfite (BS) sequencing (BS-seq) (25,38). BS-seq on mRNAs remains challenging due to high RNA degradation during the protocol resulting in low mRNA coverage (39). To increase the confidence in all discovered sites, we used two independent cell lines, the embryonic cell line Hues9 (H9) and HEK293 (HEK) cells. As negative controls, we generated two or three independent knockout clones via CRISPR/Cas9 genome editing for each cell line (Figure S2A). Furthermore, we over-expressed NSUN6 in two independent H9 clones and rescued two HEK knockout clones by stably over-expressing the NSUN6 construct (Figure S2A). At least four replicates from each condition were subjected to the BS conversion protocol, generating a total of 68 RNA BS-seq datasets. For the analyses of the RNA BS-seq data we used our established pipeline (Figure 2A; Table S3) (https://github.com/susbo/trans-bsseq) (21).

**Figure 2.**
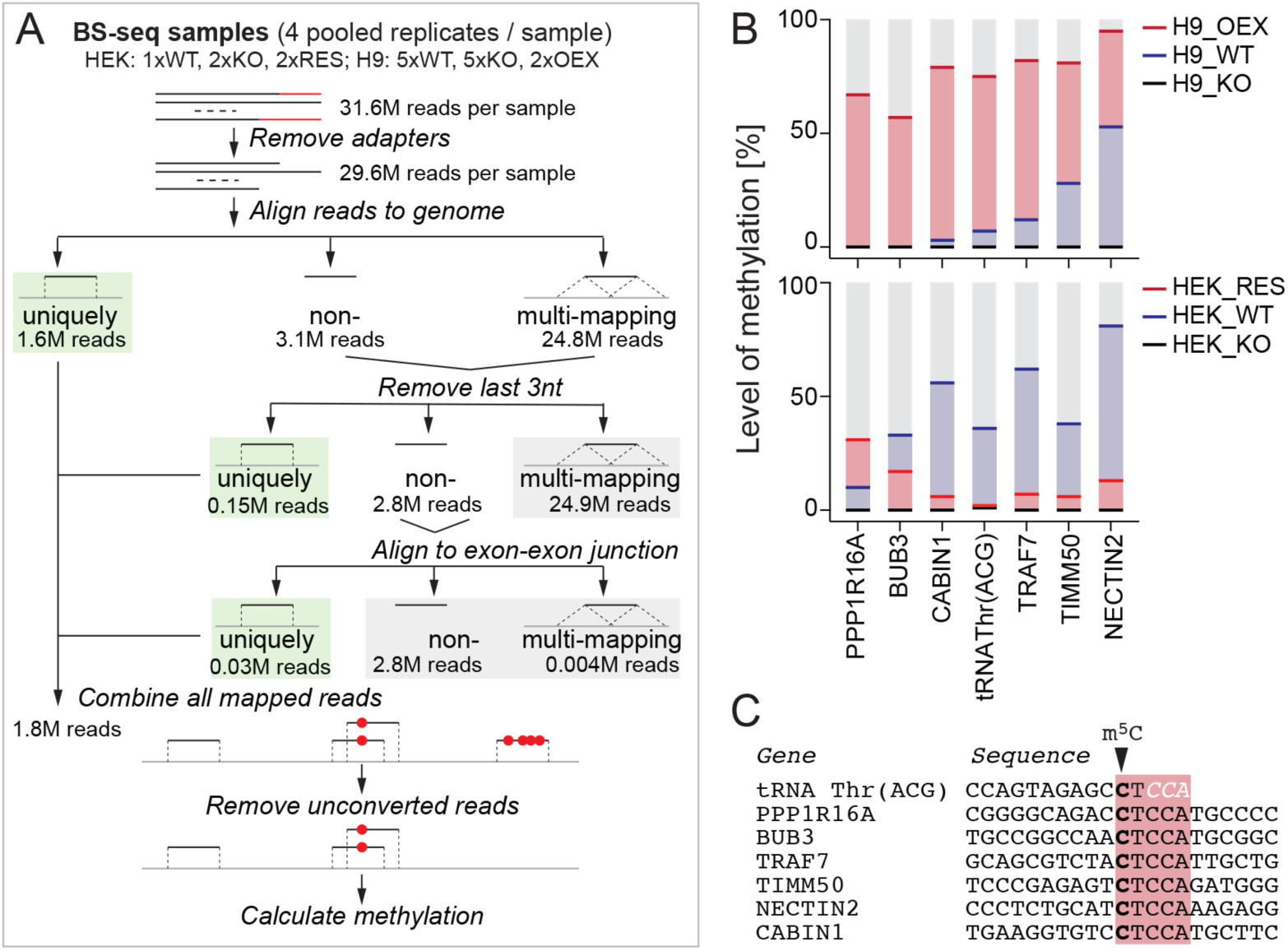
BS-sequencing confirms NSUN6 methylation sites mRNAs. (A) Pipeline of data analyses using BS-sequencing data. (B) High confidence NSUN6-dependent methylation sites identified by RNA BS-seq in H9 (upper panel) and HEK293 (lower panel). OEX: over-expression; WT: wild-type; KO: knockout; RES: rescue of NSUN6. (C) Position of the methylated site detected by RNA BS-seq and surrounding sequence. All sites contain CTCCA NSUN6 consensus motif.

After mapping the reads, all technical replicates were pooled to achieve the highest number of reads per cytosine in each condition. In summary, we analysed the following conditions: (*i*) wild-type cells with endogenous expression of NSUN6, (*ii*) NSUN6 knockout cells (Figure S2B, C), (*iii*) NSUN6 rescued cells (Figure S2D), and (*iv*) NSUN6 over-expressing cells (Figure S2E).

We first identified *de novo* NSUN6-methylated cytosines in the BS-seq datasets without using the miCLIP data as guidance. As expected, the analysis was limited by the overall low methylation levels and low mRNA read coverage at cytosines. Therefore, we pooled (*i*) all knockout (n=20), (*ii*) all over-expressing (n=8) and (*iii*) all wild-type/rescue (n=40) samples. We identified only seven high confidence NSUN6-mediated m^5^C sites in these three groups, when selecting all sites with >20 reads and <0.5% methylation in the knockout, >10 reads and >50% methylation in the over-expressing, and >30 reads and >10% methylation in the wild-type/ rescue samples (Figure 2B; Table S4). The detection of only a low number of methylated sites is in line with previous data showing that the vast majority of mRNA m^5^C sites are of low abundance, and that high abundance m^5^C sites are in fact rare in mRNA (11).

While only one of our detected sites was in a tRNA, tRNA^Thr^ (ACG) -a previously known NSUN6-target (20), all other detected m^5^C sites located to coding RNAs (Figure 2C). All of the detected sites displayed the NSUN6 target motif CTCCA, when taking the CCA-editing of tRNAs into account (Figure 2C). Five of the six m^5^C sites have also independently been previously identified as NSUN2-independent sites in HeLa cells (Table S4) (11,13).

Next, we used the RNA BS-seq data to quantify the methylation levels across all miCLIP sites within protein coding regions (Table S5A). As expected, only a small fraction of miCLIP sites were covered by more than 50 reads in the BS-seq data (Figure 3A). However, NSUN6 miCLIP sites were also consistently detected in previously published BS-seq datasets (11,13), and the greatest overlap was found in NSUN2-knockout cells (Figure 3B). Moreover, more than 80% of methylation sites showing the NSUN6 recognition motif were reduced in knockout cells and rescued or increased by over-expression of the NSUN6 construct in knockout HEK and wild-type H9 cells (Figure 3C-G; Figure S3A; Table S5B). Our finding that only CTCCA-containing methylation sites depended on NSUN6 was confirmed in previously published datasets (11,13) (Figure 3H-J). All other sites showed no difference in methylation when NSUN6 was depleted or over-expressed (Figure 3K; Table S6).

**Figure 3.**
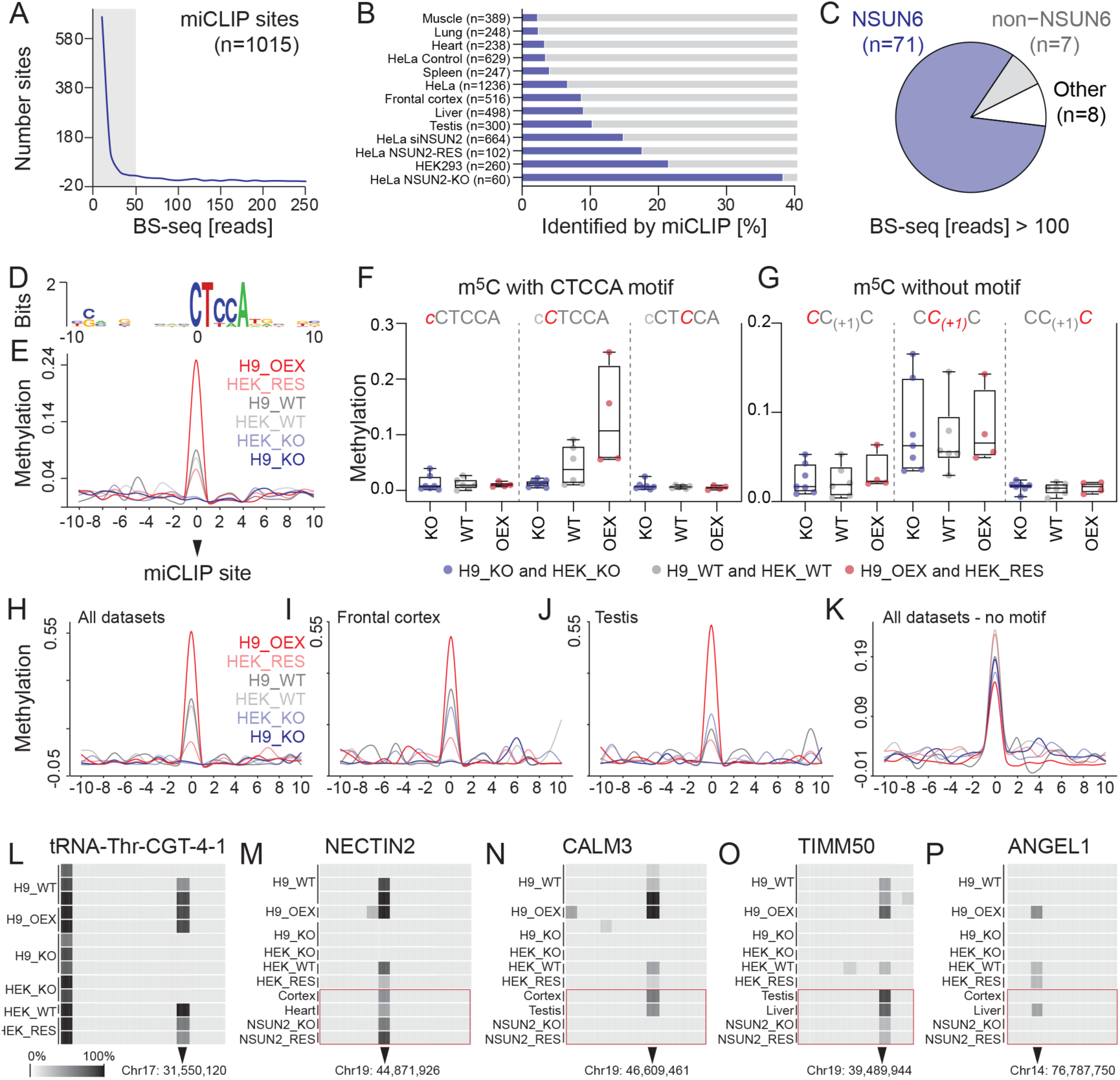
Methylation at miCLIP sites. (A) Number of protein coding miCLIP sites and coverage of BS-seq reads. Grey box: miCLIP sites covered by < 50 reads by BS-seq. (B) Fraction of methylation sites identified in the indicated datasets overlapping with NSUN6 miCLIP sites. (C) 83% of all miCLIP sites containing CTCCA motif and covered by > 100 BS-seq reads shown in (A) are NSUN6-dependent (blue). (D, E) BS-sequencing confirms consensus motif (D) at seven NSUN6-dependent methylation sites (E). (F, G) Methylation levels in knockout (KO), wild-type (WT), and NSUN6-overexpressing (OEX) cells at miCLIP sites containing the consensus motif (F) and sites without the consensus motif (G). Shown are biological replicates in H9 and HEK293 cells around the miCLIP site. (H-K) Methylation at published methylations sites (11) containing CTCCA motif identified in all datasets (H) or in selected tissues of frontal cortex (I) and testis (J) depend on NSUN6. Published methylation sites without CTCCA motif are not NSUN6-dependend (K). (L-P) Representative heatmaps showing the methylation level in the indicated samples in tRNA^Thr^ (CGT) (L) and selected mRNAs (M-P). Red box: m^5^C sites identified by Huang et al. (11) Shown are all cytosines surrounding the m^5^C site.

NSUN6 has previously been identified to methylate position 72 in tRNAs^Thr^ (TGT and AGT) and tRNA^Cys^ (GCA) (20,40), which all contain the CTCCA motif as mature tRNAs, after addition of the CCA sequence. Our data identified tRNAs^Thr^ (TGT and AGT) as NSUN6 targets (Figure 3L). tRNA^Cys^ (GCA) was not covered in our dataset and no additional tRNAs were methylated by NSUN6. All other sites located to protein coding RNAs (Figure 3M-P; Table S5A). Although the variability of methylation levels between conditions was high (Figure S3A), the identified NSUN6-dependent sites were also detected in recently published BS-seq-based transcriptome-wide screens (Figure 3M-P; red boxes) (11,13).

Mass spectrometry analysis revealed that only deletion of NSUN6 simultaneously with NSUN2 and DNMT2 resulted in a detectable reduction of m^5^C levels in tRNAs and large RNAs (Figure S3B, C), confirming that NSUN6-dependent m^5^C sites are rare, and indicating at least some enzymatic redundancy in RNA cytosine-5 methylation.

### miCLIP sites with CTCCA motif coincide with translation termination

To test whether non-methylated tRNA^Thr^ and tRNA^Cys^ caused codon biases during mRNA translation in NSUN6-depleted cells, we performed ribosome profiling. Codons for threonine and cysteine were neither enriched nor depleted the ribosome active sites; however, we found stop codons to be significantly under-represented at the ribosomal A site in NSUN6 knockout cells (Figure S4A). Since the process of translation termination begins when the ribosome encounters a stop codon in the ribosomal A site (41,42), we next asked how translation was affected at the miCLIP sites.

To analyse mRNA translation, we considered all miCLIP sites in protein coding regions containing the CTCCA motif (n=994) and calculated the fraction of annotated coding sequences within 3000 bases centred on the motif (Figure 4A). We measured a higher frequency of coding sequences up-stream than down-stream of the miCLIP sites, indicating that miCLIP sites more often coincided with a stop of translation (Figure 4A). To investigate this further, we divided the miCLIP sites into three groups based on their Ribo-seq coverage, now marking the start (more reads down-than up-stream), middle (equal read distribution up- and down-stream) or end (more reads up-than down-stream) of translation (Figure 4B-D; upper panels). The CTCCA motif coincided with translation termination at the majority of miCLIP sites (Figure 4D; lower panel), which was further illustrated by the sharp drop in translation across those miCLIP sites (Figure 4E). The remaining miCLIP sites marked the start codon or were evenly distributed (Figure S4B, C). In contrast, RNA-seq reads distributed both up- and down-stream of miCLIP sites suggesting that NSUN6-binding sites marked the end of translation but not end of the corresponding mRNA (Figure 4F; Figure S4D, E). Thus, our analysis revealed that the putative methylated sites marked translation termination.

**Figure 4.**
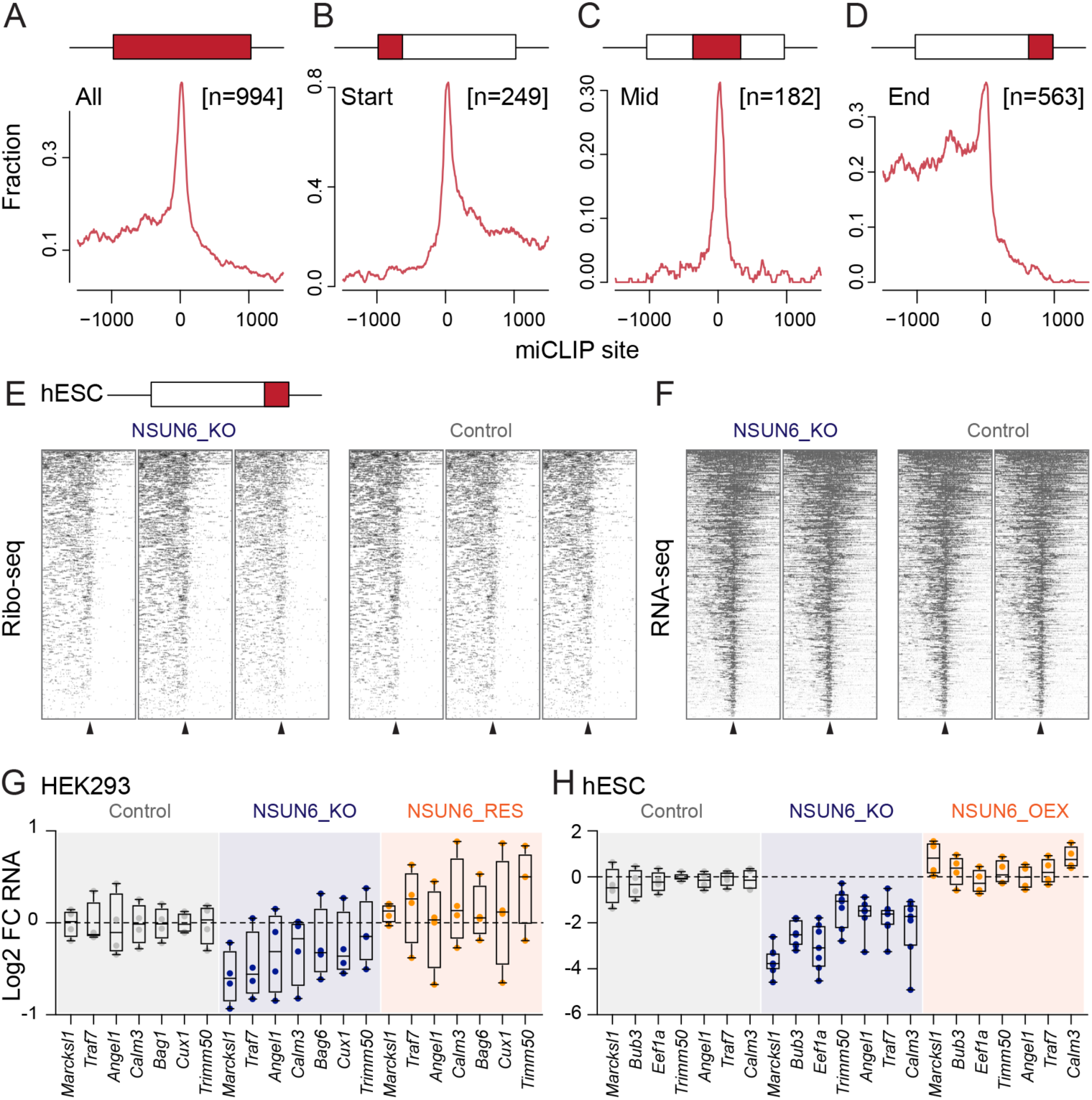
NSUN6 miCLIP sites with CTCCA consensus motif mark translation termination. (A-D) Annotated coding sequence 1500 nucleotides up- and down-stream miCLIP sites containing the CTCCA consensus motif using all NSUN6 miCLIP sites (n=994) (A), or only sites classified as the start (n=249) (B), middle (n=182) (C), or end (n=563) of translation. (E, F) Heatmaps showing the Ribo-seq (E) or RNA-seq (F) read coverage in NSUN6 knockout clones (H9) (left panels) or controls (right panels). Arrowheads indicate position 0 of miCLIP sites. (G, H) RT-QPCR for selected mRNAs carrying NSUN6-methylated cytosine in HEK cells (G) and the human embryonic stem cell line H9 (hESCs) (H) in control, knockout (KO), over-expressing (OEX), or rescued (RES) cells. Data are normalized to 18s rRNA and shown relative to control cells. Outlier (highest and lowest values) are removed in (H).

### Loss of NSUN6-mediated methylation decreases mRNA stability

Since CTCCA motif containing miCLIP sites predominantly occurred in 3’UTRs (Figure 1F), we concluded that ribosomes either had restricted access to the 3’UTR or were removed to ensure translation termination. As mRNA stability and translation are intertwined (43), we asked whether RNA stability of NSUN6-targeted mRNAs was affected by removal of NSUN6. Indeed, RNAs with exonic miCLIP sites were significantly less abundant in NSUN6-depleted cells (Figure S4F). To confirm that NSUN6-targeted mRNAs were less stable in absence of NSUN6, we performed RT-QPCR (Figure 4G, H). We measured a reduction of mRNAs with confirmed methylated sites in both NSUN6-depleted cell lines, which were rescued by re-expressing or over-expressing the wild-type NSUN6 protein (Figure 4G, H). Thus, loss of methylation at CTCCA motifs slightly decreased mRNA level, potentially caused by lower translation.

However, the RNA sequencing data failed to reveal a consistent set of transcripts affected by removal of NSUN6. Since RNA modification enzymes often act directly in response to external stimuli (2), we asked whether differentiation of human embryonic stem cells was affected when NSUN6 was depleted. We differentiated the H9 cells into embryoid bodies and confirmed reduced levels of NSUN6-targeted mRNAs in embryoid bodies lacking NSUN6 (Figure S5A). While pluripotency factors were not differentially expressed, some mesoderm markers and *Hoxa1* were consistently reduced in the absence of NSUN6 (Figure 5A). Finally, we asked whether loss of NSUN6 affected embryonic development. To generate total knockout mice, we used two embryonic stem cells clones carrying the LacZ-reporter in exon 2 of the *Nsun6* gene leading to transcriptional disruption of a functional NSUN6 protein (Figure S5B). Although, LacZ expression revealed that *Nsun6* was quite ubiquitously expressed, we observed no gross phenotype in the absence of NSUN6 (Figure 5B-M). We concluded that embryonic development was unaffected by total loss of NSUN6.

**Figure 5.**
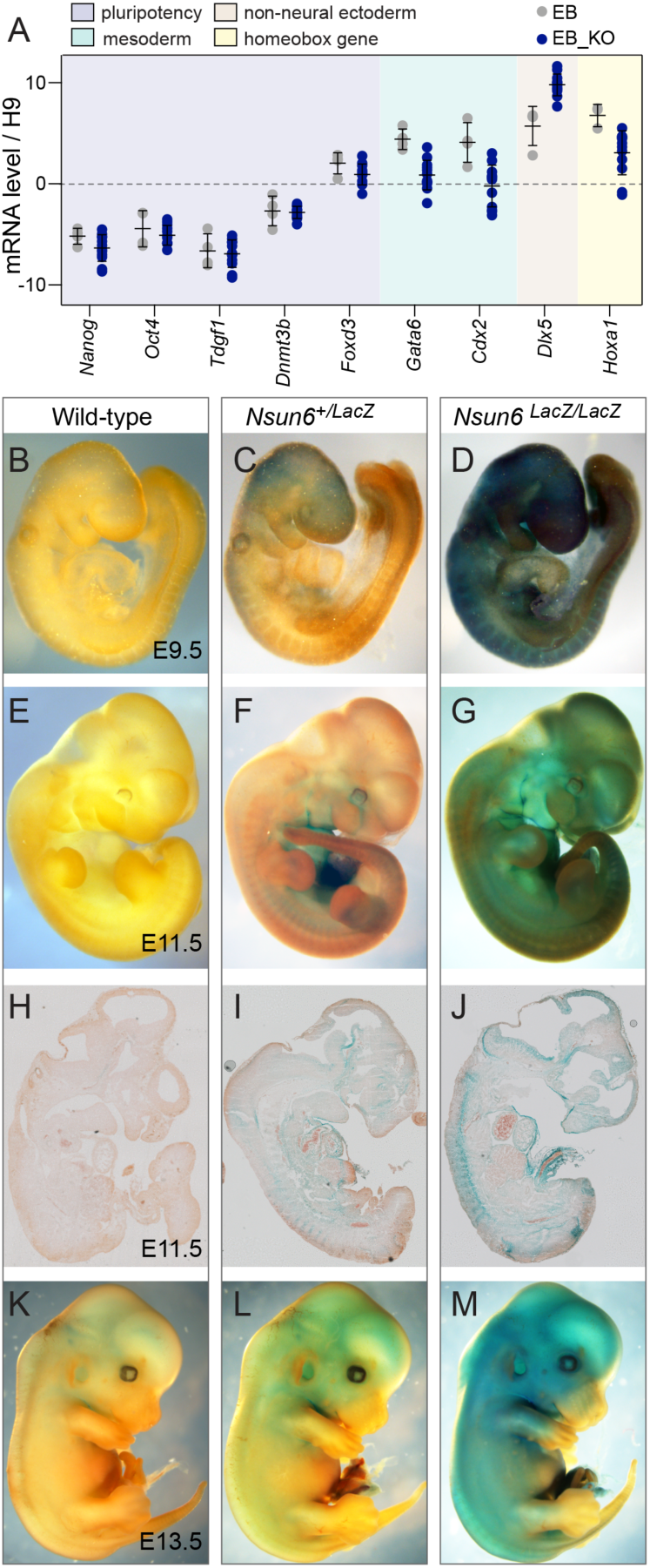
NSUN6 is dispensable for embryonic development. (A) RT-QPCR for RNA of markers for pluripotency, mesoderm, non-neural ectoderm, and a HOX gene in embryoid bodies expressing (EB) or lacking NSUN6 (EB_KO). Data are normalized to 18s rRNA and shown relative to self-renewing H9 cells. (B-M) Mouse embryos at the indicated embryonic (E) days are labelled for LacZ expression. Images are showing the whole embryo (B-G, K-M) or histology of a section (H-J).

In summary, our data identify NSUN6 as a methyltransferase targeting mRNAs at a CTCCA motif near the translation termination sites primarily in 3’ UTRs. Although loss of NSUN6-mediated methylation decreased mRNA stability of some targets, NSUN6 is not required for embryonic development.

## DISCUSSION

RNA modifications add flexibility, diversity and complexity to cell type and state-specific gene expression programs. Mapping these RNA modifications on single nucleotide resolution is a critical step towards understanding the underlying regulatory pathways. Despite recent advances in technologies to detect RNA modifications in mRNAs (3,44-46), the prevalence and precise location of in particular rare modifications remain often unclear (9). Although the need for more robust detection methods was recognised early on in the field (47), RNA BS-seq remains the method of choice for mapping m^5^C transcriptome-wide because it is currently the only available method detecting m^5^C sites on endogenous mRNAs in a quantitative manner (39).

Optimizing RNA BS-seq protocols and computational analyses is one strategy to improve the detection of m^5^C (8,11). The most recent study identified about 100 m^5^C sites per megabase in a given mammalian tissue or cell type (11). Huang et al (11) further demonstrated that many but not all detected sites were NSUN2-dependent, and that NSUN2-independent sites were marked by a 3’TCCA motif. The authors suggested that this sequence motif was targeted by an uncharacterised cytosine-5 RNA methyltransferase (11). In our study, we discover the same consensus motif (CTCCA) in mRNAs and further demonstrate that it is in fact targeted by the characterised cytosine-5 methyltransferase NSUN6. Furthermore, the known tRNA targets of NSUN6 also carry this motif, but only after addition of the CCA tail as mature tRNA (20).

The identification of NSUN6 as an additional methyltransferase that can target mRNA might explain why the prevalence of m^5^C has been controversial. Potential redundancy of these methyltransferase towards mRNA may hamper studying the functional relevance of m^5^C and explain the apparent lack of phenotype in NSUN6-depleted mice. However, we find that NSUN6-targeted consensus motifs coincide with translation termination and the stability of targeted mRNAs were reduced in the absence of NSUN6. Our finding that candidate target mRNAs are more stable in the presence of m^5^C is in line with the recent finding that m^5^C protects maternal mRNA from decay in the maternal-to-Zygotic transition (48).

Together, our investigations confirm that m^5^C in a transcriptome is relatively rare but can at least in some cell and tissue types be mediated by two methyltransferases, NSUN2 and NSUN6.

## CODE AVAILABILITY

The scripts used for the alignment and processing of the BS-seq data are available at [https://github.com/susbo/trans-bsseq].

## DATA AVAILABILITY

NGS data are available on GEO (GSE125046). RNA-seq, Ribo-seq, miCLIP data are up-loaded onto GEO (GSE140997)

## ACKNOWLEDGMENTS

We thank everybody who provided us with reagents. We gratefully acknowledge the support of all the Wellcome MRC Cambridge Stem Cell Institute core facility managers, in particular Maike Paramor. This work was funded by Cancer Research UK (CR-UK), the Medical Research Council (MRC) and the European Research Council (ERC). Parts of this research in Michaela Frye’s laboratory was supported by core funding from Wellcome and MRC to the Wellcome MRC Cambridge Stem Cell Institute.

---

**Figure S1.**
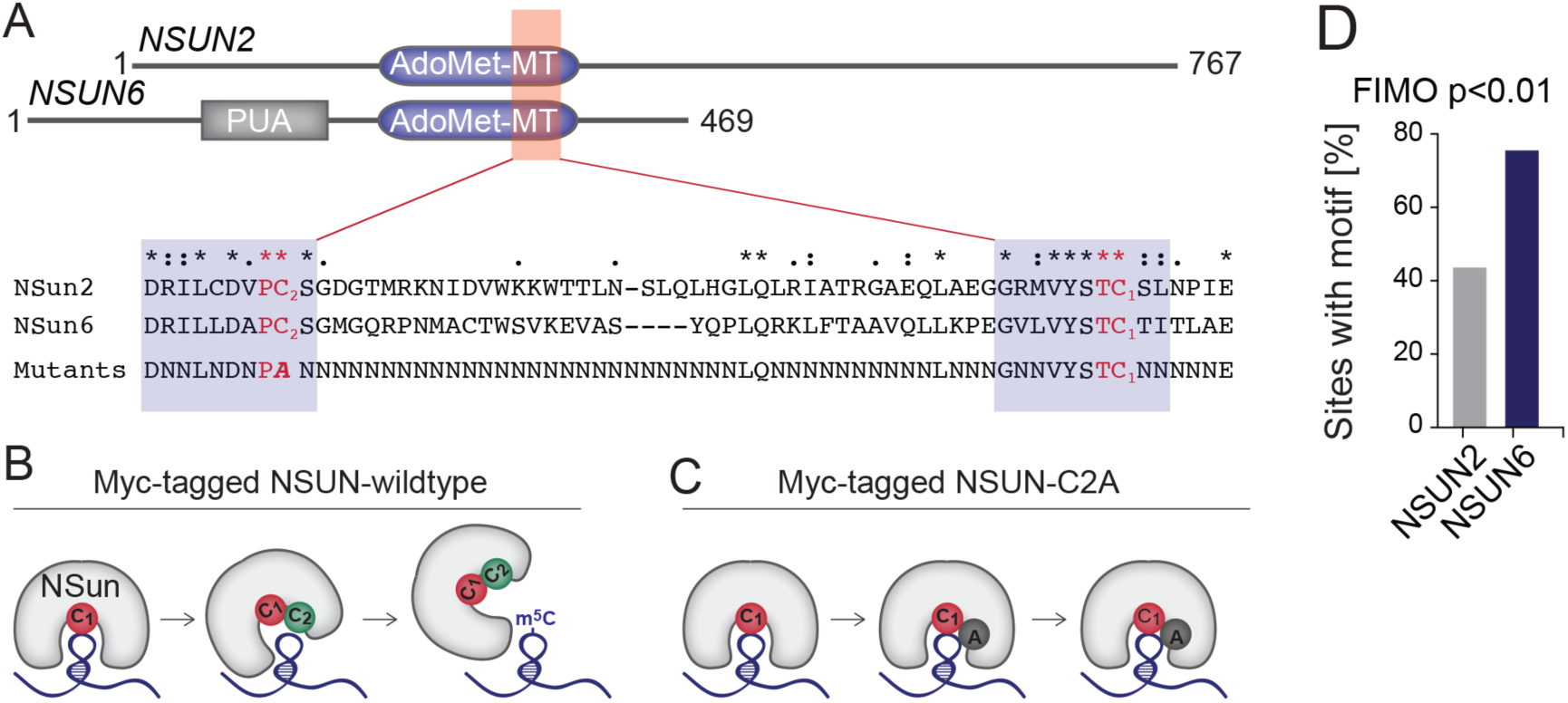
MiCLIP identifies NSUN6 targeted RNA. (A) Schematic overview of conserved protein domains in NSUN2 and NSUN6 showing the Ado-Met methyltransferase (MT) superfamily domain and the amino acid sequence around the MT domain. Red letters indicate the catalytic active sites and the mutation to generate the miCLIP constructs (bold). NSUN6 contains an additional pseudouridine synthase and archaeosine transglycosylase (PUA)-domain, a conserved RNA-binding domain. (B,C) Overview of the mutated Myc-tagged NSUN-constructs and how they form m^5^C (B) or the covalent bond at the methylated site (C). (D) Percentage of NSUN2 and NSUN6 miCLIP sites showing the consensus sequence motif shown in Figure 1H, I.

**Figure S2.**
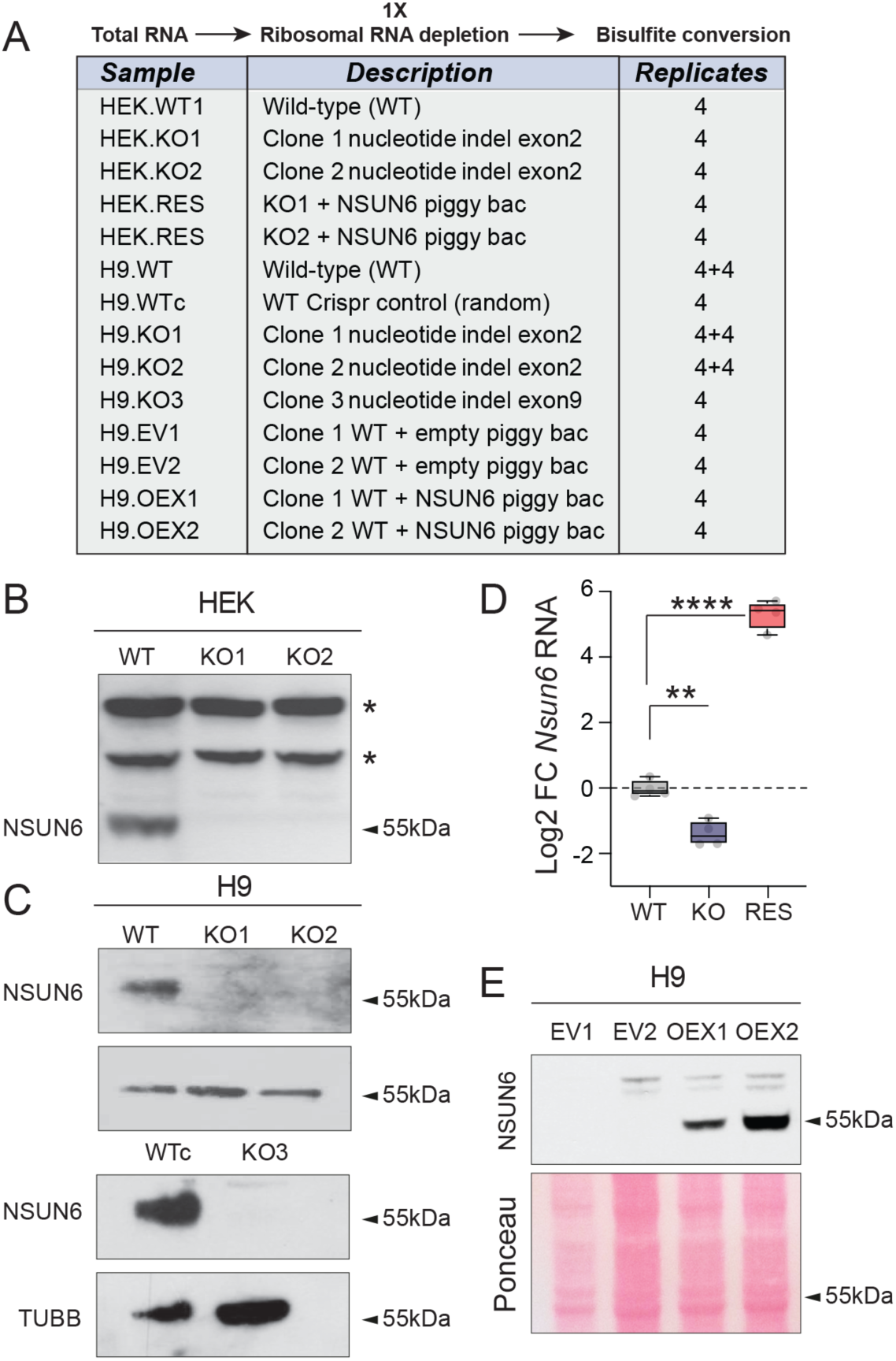
Sample generation and detection of m^5^C by mass spectrometry. (A) Preparation for bisulfite conversion protocol and list of all samples analysed using RNA BS-seq. (B) Western blot showing loss of the NSUN6 protein in HEK cell clones. Asterisks: non-specific bands serving as internal loading controls. (C) RT-QPCR for *Nsun6* RNA in knockout (KO) and rescued (RES) HEK cells. (D) Western blot showing loss of the NSUN6 protein in three H9 cell clones (KO1-3). WTc: H9 clone with random integration of an indel. Tubulin (TUBB) served as loading control (B, D). (E) Western blot showing over-expression (OEX) of the NSUN6 protein in H9 cells. EV: Empty vector. Ponceau labelling of proteins served as loading control.

**Figure S3.**
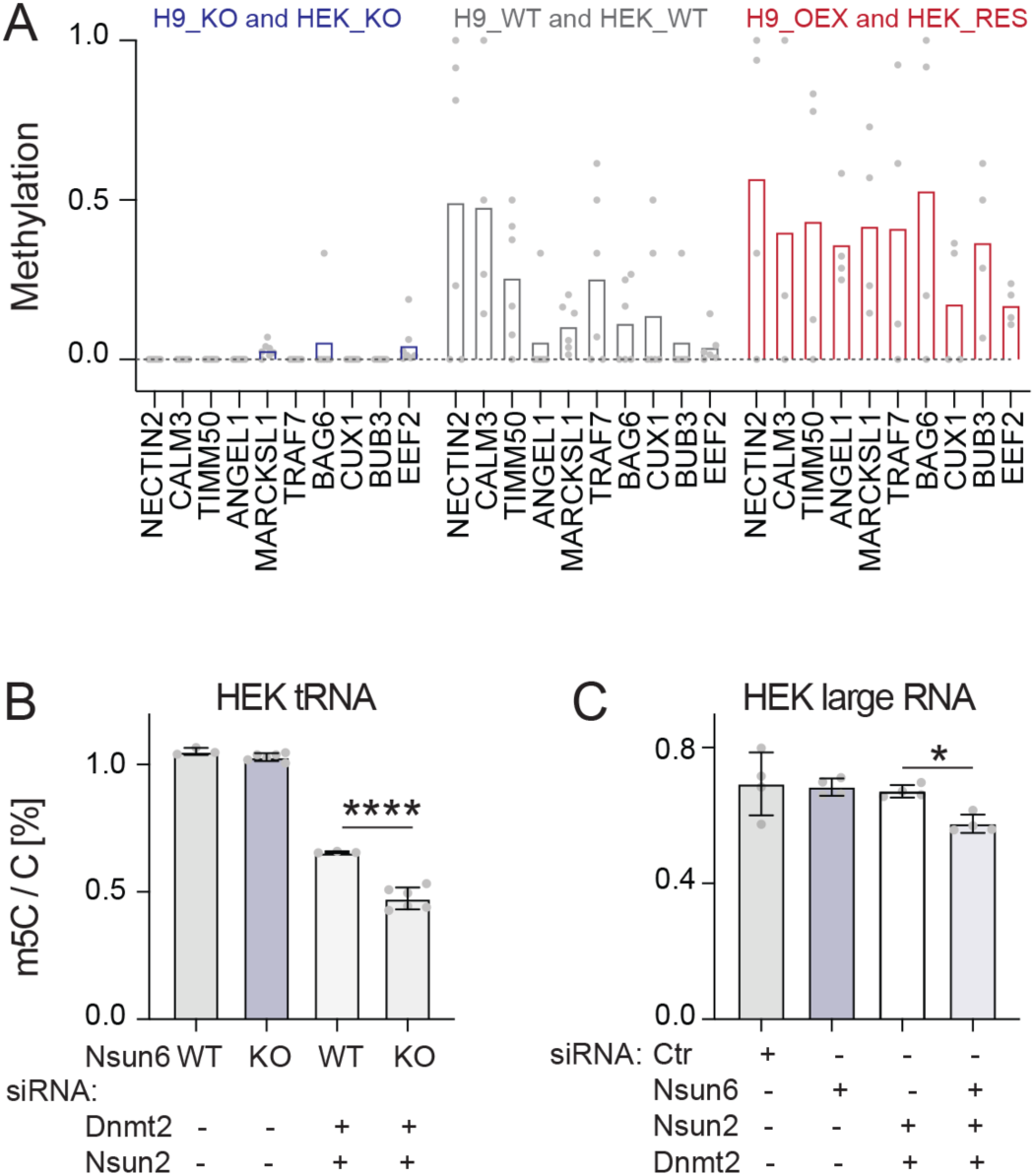
RNA bisulfite sequencing and mass spectrometry to detect m^5^C. (A) Methylation level at miCLIP sites containing the CTCCA motif in wild-type (WT) H9 and HEK cells, NSUN6 knockout (KO) cells, and NSUN6 rescued (RES) or cells over-expressing (OEX) cells. Bar plots indicate the mean methylation level in all samples. (B, C) Mass spectrometry measuring m^5^C in HEK cells enriched for tRNAs (B) or large RNAs (C) using HEK wild-type (WT) and NSUN6 knockout (KO) (B) or cells transfected with the indicated siRNAs (C).

**Figure S4.**
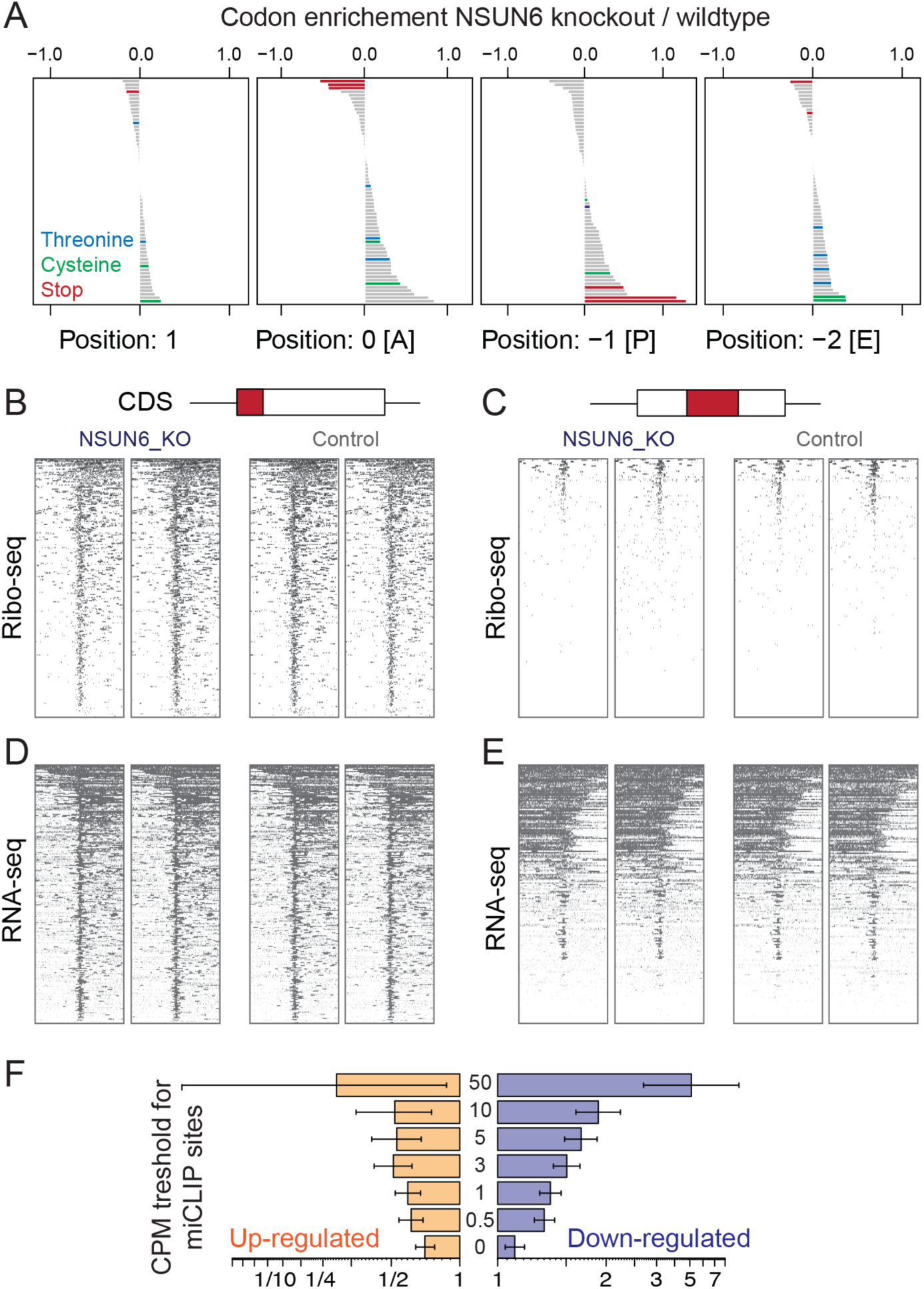
Ribo-seq and RNA-seq analyses in human embryonic stem cells and HEK cells. (A) Codon enrichment analyses at the ribosome position 1 and the A, P, and E sites in wild-type and NSUN6-depleted human embryonic stem cells (H9). (B-E) Distribution and positional profiles of Ribo-seq (B, C) and RNA-seq (D, E) up- and down-stream miCLIP sites at start (B, D) or middle of translation (C, E). (F) Odds ratios describing a higher occurrence of down-regulation (blue) and lower occurrence of up-regulation (red) in genes with exonic miCLIP sites compared with expressed genes without miCLIP sites, at the indicated CPM thresholds. Odds ratios and 95% confidence intervals were calculated using Fisher’s exact test.

**Figure S5.**
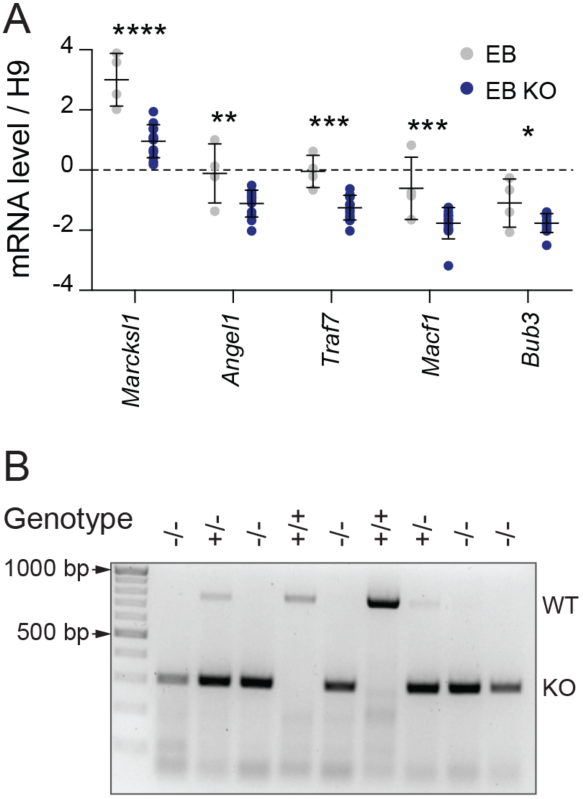
NSUN6 target mRNAs are less stable in the absence of NSUN6. (A) RT-QPCR for RNA for mRNAs with confirmed methylation sites in wild-type embryoid bodies (EB) and NSUN6-depleted EB. Data are normalized to 18s rRNA and shown relative to self-renewing H9 cells. (B) Representative PCR for genotyping *Nsun6* -/-, +/-, and +/+ mice.

